# Cisplatin and Copper Induce Distinct Endocytic Pathways of Copper Transporter-1 and Elicits Unique Response on Copper Homeostasis Proteins

**DOI:** 10.1101/2025.11.03.686439

**Authors:** Mainak Chakraborty, Asmita Pal, Sumanta Kar, Arnab Gupta

## Abstract

Cisplatin (CDDP), a widely used chemotherapy drug is imported in mammalian cells by copper transporter CTR1. We observed that CDDP cytotoxicity is significantly reduced in CTR1 deficient yeast and in copper loaded mammalian cells indicating its shared uptake pathway with copper. CTR1 resides on the plasma membrane undergoes endocytosis during CDDP import as in copper, involving its amino-terminal His-Met-motifs and the conserved ^150^MXXM^154^ pore motif. Unlike Copper, CDDP directs endocytosed CTR1 toward lysosomes rather than being sorted through VPS35-positive endosome. CDDP enhances lysosomal acidity and cathepsin activity. Furthermore, CDDP selectively downregulate cuproproteins, CCS and SOD1, disrupting copper homeostasis. This study highlights the unique aspects of copper versus CDDP uptake by CTR1 and its differential downstream ramification on lysosomes and cuproproteins.

## Introduction

Transition metals, manganese, iron, cobalt, copper, zinc are essential micronutrients that play pivotal roles in regulation of cellular homeostasis in all eukaryotic life forms (1). Copper is one of the *d*-block transition metals known to act as cofactor for wide range of metalloenzymes in cells (2). Being a member of transition metals, copper shows two different oxidation states, Copper(I) and Copper (II). In general Copper(I) is the biologically active form and Copper (II) to Copper(I) reduction at cell surface is crucial before uptake in the cell, as revealed from studies in yeasts and human copper import (3–5). Copper’s has the potential to undergo Fenton-Like reaction and generate highly toxic redox active molecules, that causes damage to the cell (6). Copper deficiency and copper excess in the body lead to neurological and hepatic disorders like Menkes disease and Wilson disease respectively (7). To protect the cells from high Copper toxicity, organisms have developed intricate machinery involving several proteins to maintain the level of bioavailable copper. Copper, after entering the cell through the Copper Transporter-1, CTR1, is subsequently sequestered by copper chaperones in the cytosol. Excess copper is then exported from the cell via copper ATPases ATP7A/B. This coordinated mechanism helps cell to maintain proper copper homeostasis (6–10)

CTR1(*SLC31A1*) is the sole high affinity copper importer, evolutionarily conserved in eukaryotes from invertebrates to higher order chordates and ubiquitously expressed in all human cells (11,12). Human CTR1, hCTR1 is an integral membrane protein consisting of 190 amino acid residues. It features a long extracellular N-terminal region rich in charged amino acids, three transmembrane segments, and a short 12-residue C-terminal tail, and that terminates with conserved HCH motifs (13,14). Ctr1 imports copper in a unique manner, distinct from traditional ion channels. Its function depends on a trimeric conformation, which is essential for its activity. X-ray crystallographic studies of *Salmo Salar* Ctr1 revealed that the transporter functions as a homotrimer, selectively importing copper through a trans-chelation mechanism. This structural insight underscores the unique mechanism by which Ctr1 facilitates copper uptake (15). This process is facilitated by charged amino acid residues, which involve two distinct clusters of methionine triads on the extracellular side and cysteine residues on the cytosolic terminal. These residues are crucial for coordinating copper ions, ensuring efficient transport. Once copper is imported, it is taken up by copper chaperones for intracellular distribution (16,17). Ctr1 is found both in the plasma membrane and in intracellular vesicles, depending on the cell’s copper levels. It maintains its abundance at the plasma membrane through a self-regulatory copper-sensing mechanism, and is internalized when extracellular copper levels exceed a threshold, thereby preventing copper toxicity (18). The internalized Ctr1 population exhibits distinct vesicular localization. Ctr1 follows a clathrin-mediated endocytic pathway, initially accumulating in early sorting endosomes, which are marked by EEA1 (19). Clifford et al. also demonstrated that upon copper depletion, Ctr1 can be recycled back to the plasma membrane via two distinct routes: a fast-recycling pathway mediated by Rab4 GTPase and a slower recycling route marked by Rab11 GTPase (20,21). Additionally, the Cullen group established the role of the retromer complex in the regulated retrieval of Ctr1 from recycling compartments (22).

Ishida et al. demonstrated that, besides copper uptake, Ctr1 is responsible for the accumulation of the anticancer drug cisplatin, observing that deletion of Ctr1 in yeast led to cisplatin resistance (23). Studies from Howell group in Human ovarian carcinoma cells also supported pharmacological importance of Ctr1 in cisplatin uptake (24,25). Cisplatin (CDDP; *cis*-[Pt(NH_3_)_2_Cl_2_]) received clinical approval as a chemotherapeutic drug in 1978 and has since become one of the most widely used treatments for ovarian cancer. However, over time, the efficacy of cisplatin has diminished due to the development of resistance, leading to therapeutic failure and increased mortality rates (26,27). Lung and ovarian cancer Patients sample analysis indicates increased in expression of copper efflux transporters ATP7A and ATP7B during Cisplatin treatment may induce drug resistance (28). Ctr1 also plays a role in promoting cisplatin resistance. Contradictory results of Ctr1 expression have been observed across different cell lines. While previous findings from ovarian and cervical cancer cell models stipulated significant degradation of Ctr1 upon treated with Cisplatin thus generate drug resistance, Kalayda et al observed no change in levels of Ctr1 (24,29,30).

The fate of the plasma membrane population of Ctr1 in the presence of copper is well established (5,21,31). As mentioned earlier, methionine and Histidine residues in Ctr1 play a crucial role in copper uptake and delivery. Under high copper conditions, Ctr1 undergoes endocytosis, localizing on early endosomes. However, the mechanism by which cisplatin enters the cell and its impact on Ctr1 endocytosis and trafficking remains poorly understood. Study by Guo et al was the first to investigate whether cisplatin can mimic copper in the regulation of Ctr1 (32). They proposed that cisplatin treatment can stabilize a high molecular weight complex of Ctr1, highlighting a unique interaction between cisplatin and Ctr1 that differs from copper’s effects (32). Using FRET assays, another study reported that Ctr1 complex formation upon copper treatment is distinct from Cisplatin treatment (33).

In this study, we aim to determine the impact of copper and cisplatin on the divergent fate of Ctr1. We mapped and compared the different endosomal localizations of Ctr1 upon copper and cisplatin treatment. Utilizing mutational analysis, we identify the specific amino acid residues of Ctr1 responsible for cisplatin import, in contrast to copper. Finally, we determined the effect of cisplatin on metallochaperones that bind copper.

## Results

### CTR1 facilitates CDDP uptake in cancerous and non-cancerous cell lines

Cisplatin or CDDP has been reported to involve copper transporter, CTR1 for its uptake; however, the reports have often been debatable. Cisplatin, exhibits multiple levels of cellular response following its entry into cells (23,34).We determined CDDP-induced cytotoxicity in cancerous (HeLa) and a non-cancerous cell line (HEK293T). We tested whether CTR1 mediated copper and CDDP uptake exhibit similar uptake mechanism. CDDP treatment was competed with copper. Using the MTT assay as a measure of cytotoxicity imparted by CDDP, we observed that in basal conditions or in absence of copper (i.e., treated by copper chelator TTM), survival of HeLa and HEK293T cells falls rapidly as compared to copper pretreated cells (**Fig. 1A, 1B**). Noticeably, CDDP is able to exert higher cytotoxicity in absence of copper and is unable to do the same when Copper is abundantly loaded in cell. This signifies the involvement of copper uptake mechanism in CDDP-induced cytotoxic.

**Fig. 1.**
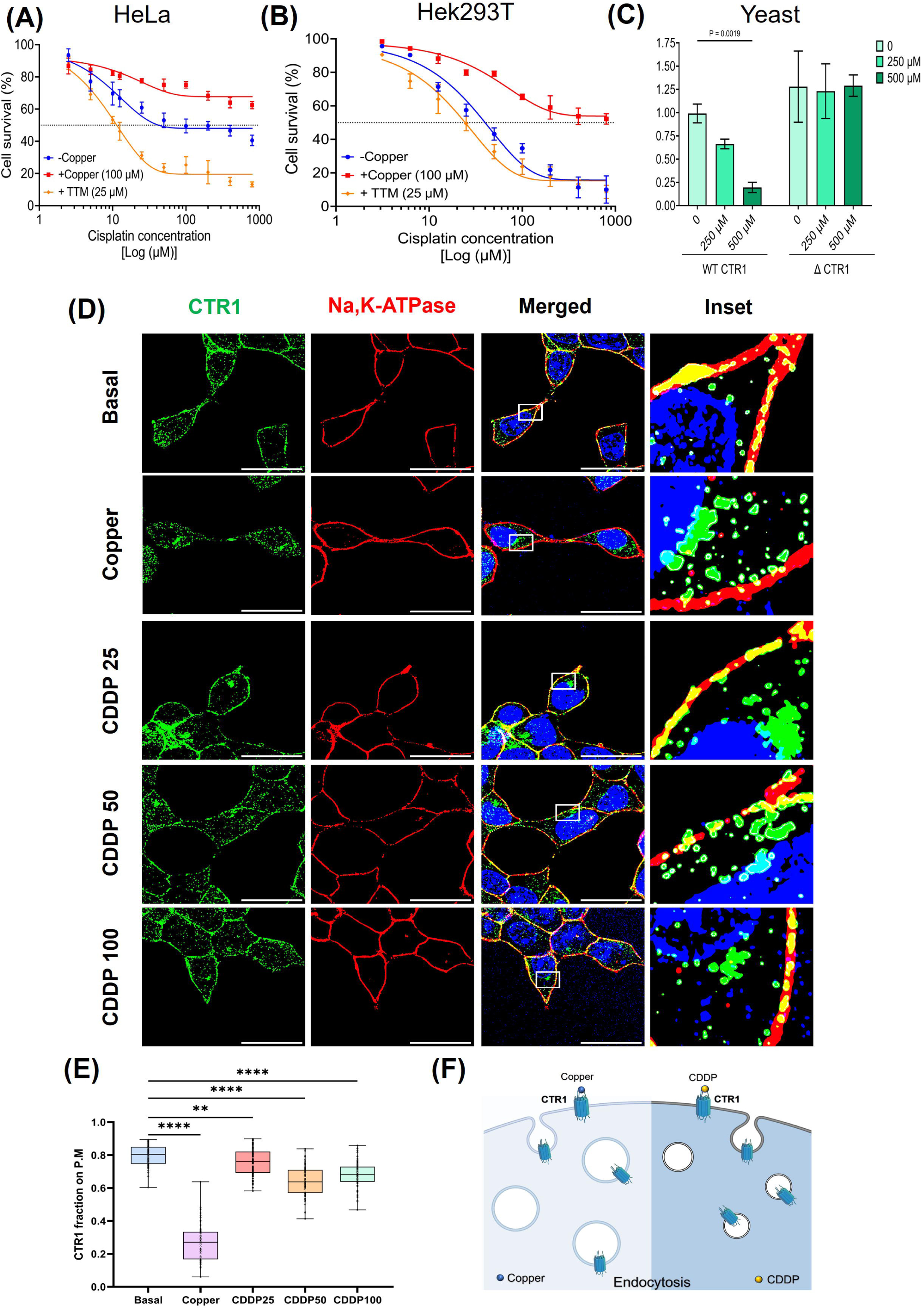
Cell survival assay and endocytosis of CTR1 in response to Copper and CDDP Cell viability of. **(A)** Cancerous cell line HeLa and **(B)** Non-cancerous cell line Hek293T when pre-treated with copper (100 µM) or copper chelator TTM (25 µM) for 1 hour followed by different conc. of CDDP. CDDP concentration (µM) has been depicted in logarithmic units in x-axis. Non-linear regression curve analysis has been carried out. The dashed line through the center represents 50% cell inhibition. (**D)** Growth curve of yeast (*S*. *cerevisiae*) BY4742 strain WT and ΔCTR1 mutants in presence of increasing doses of CDDP (250 and 500 µM). Data represent mean ± SD (n=6). **(D)** Immunofluorescence studies in HEK293T non-cancer cells shows colocalization of CTR1 (green) with Plasma Membrane marked with Na^+^/K^+^ ATPase (Red) in basal condition, get endocytosed upon administration of both copper (100 µM) and gradual increase in conc. of CDDP (25 µM, 50 µM, 100 µM) showing vesicular faction of CTR1 in the cytosol. In the merged figures DAPI (BLUE) staining indicates nucleus and the marked area on ‘merge’ panel is enlarged in ‘inset’. The arrowhead shows colocalization between Na^+^/K^+^ ATPase – Ctr1.The Scalebar represents 10 microns. **(E)** The fraction of hCTR1 colocalize with membrane marker Na^+^/K^+^ ATPase demonstrated by a Box and Whisker plot with jitter points. (n>50) The box represents the 25 to 75th percentiles, and the median in the middle. The whiskers show the data points within the range of 1.5 × interquartile range (IQR) from the first and third quartile. ∗*p* < 0.05, ∗∗∗∗*p* < 0.0001; ns, not significant (nonparametric Mann–Whitney U test/Wilcoxon rank-sum test). Statistical significance is shown above the bars measured through One-way ANOVA followed by Dunnett’s and Sidak’s multiple comparison analysis (p<0.01). **(F)** Graphical scheme representing endocytosis of Ctr1 in presence of copper and CDDP.

We further tested CDDP mediated toxicity in wild-type and yeast lacking CTR1, ΔCTR1 *Saccharomyces cerevisiae*. Corroborating with our mammalian cell findings, ΔCTR1 show no signs of CDDP cytotoxicity as compared to the yeast cells expressing WT-CTR1, where CDDP exert pronounced cytotoxicity in a dose-dependent manner. Thus, we can infer that CTR1 is the primary importer of CDDP and absence of CTR1 prevents CDDP from exerting cytotoxicity in yeast (**Fig. 1C**).

Studies from our group and others have shown that CTR1 localizes on the plasma membrane, colocalizing with the plasma membrane marker Na-K-ATPase (5). CTR1 undergoes endocytosis as a self-regulatory response upon increasing copper treatment (5). We asked whether CTR1 exhibits similar endocytic phenomenon in response to CDDP treatment. HEK293 cells were transfected with 3X-Flag-CTR1 and localization were studied in fixed cell under confocal microscope. Though to a much lesser extent, CTR1 undergoes endocytosis in response to CDDP treatment, similar to as observed in case of copper treatment (**Fig. 1D, 1E)**. Our finding corroborates with previous reports that CDDP uptake involves the copper uptake machinery and CDDP induces CTR1 endocytosis (35). A comparable response was observed with increasing drug concentrations (25 µM, 50 µM, 100 µM) induced a dose-dependent reduction in CTR1 co-localization with Na⁺/K⁺-ATPase at the plasma membrane and promoted its endocytosis into intracellular vesicles. Interestingly, the CTR1 endocytic vesicles formed by CDDP induction was appreciably smaller than those induced by copper. With increase in CDDP concentration, we noticed a slight but statistically significant increase in CTR1 endocytosis. (**Fig. 1D, 1E**)

To summarize we report that presence or absence of copper can interfere with CDDP cytotoxicity in both cancerous and non-cancerous cells. Additionally, our study shows that CTR1 endocytosis is a part of the initial signalling cascade triggered by CDDP similar to that of copper. A schematic summarizing the comparable physiological responses elicited by copper and CDDP, leading to the endocytic internalization of CTR1 from the plasma membrane, is presented in **Fig. 1F**.

### The extracellular amino-terminus and Met-motifs of the central ion conduction core of CTR1 is key for CDDP uptake

CTR1 has an extracellular amino terminal domain rich in His-Met clusters. Puig et al demonstrated the key role of conserved methionine residues in high-affinity copper uptake, through biochemical and genetic analyses of yeast and human copper transporters (36). The extracellular amino-terminus of CTR1 harbours two methionine clusters, i.e., ^7^MGM^9^ (termed here as M1) and ^40^MMMMPM^45^ (M2) and a His cluster ^3^HSHH^6^. We have previously shown that these methionine motifs regulate Copper(II) to Copper(I) reduction and regulates copper import (5). We further found that the two clusters exhibit functional complementarity, as the second cluster is sufficient to preserve copper-induced CTR1 endocytosis upon complete deletion of the first cluster (5).

To investigate the role of N-terminal amino acids of Ctr1 in CDDP uptake we used N terminal Δ^3^HSHH^6^, ΔM1(^7^MGM^9^), ΔM2(^40^MMMMPM^45^) & Δ30 mutant constructs and compared them with the WT-CTR1. The Δ30-CTR1 mutant lacks the first 30 amino acids that harbour the M1 and the His cluster (**Fig.2A**). The mutant proteins were stable and localized at the plasma membrane. We observed clear endocytosis of WT-Flag Ctr1 in vesicular compartments in response of both Copper and CDDP **Fig.2B**.

**Fig. 2.**
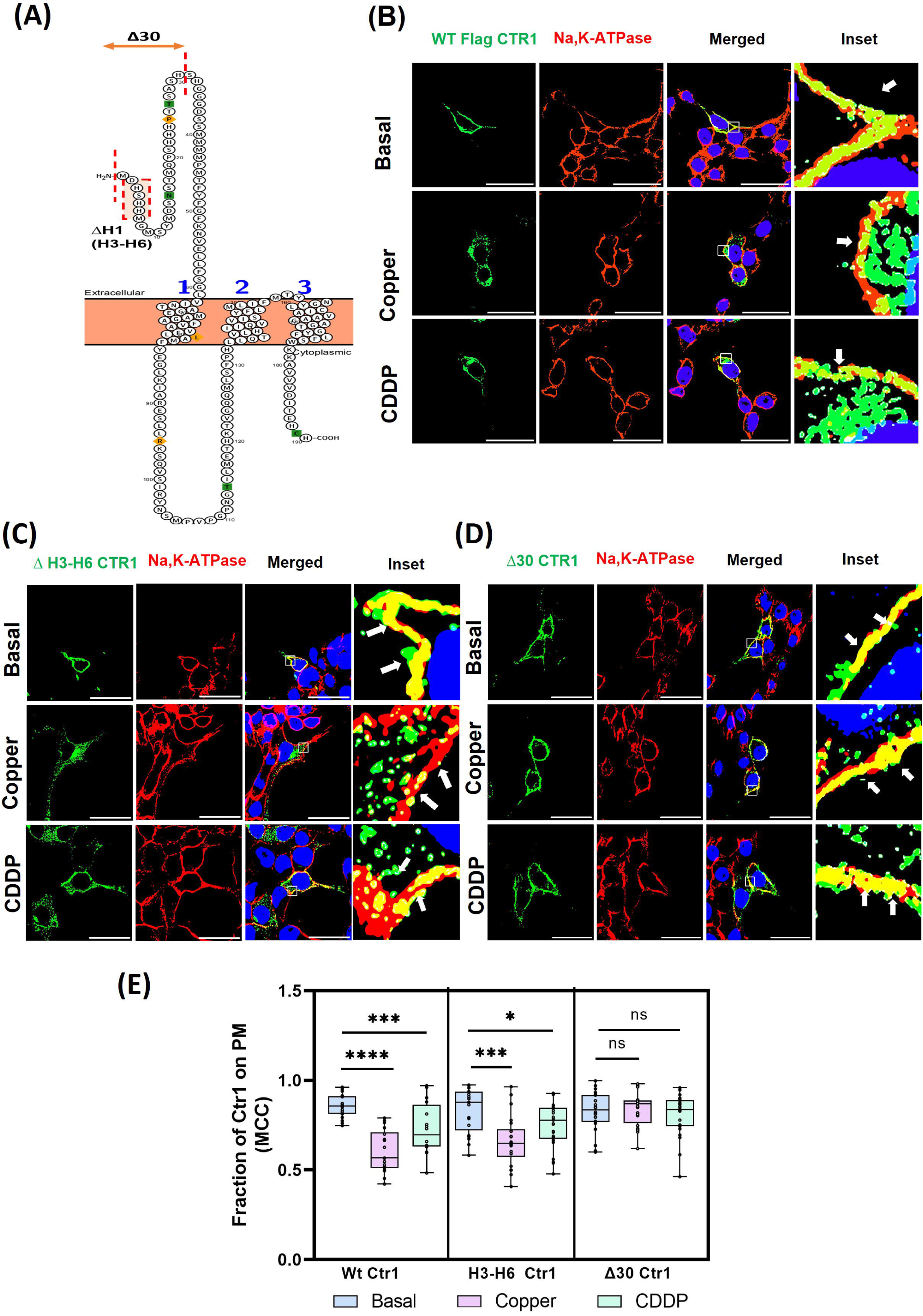
Fate of plasma membrane localized Flag-WT-Ctr1 and its amino-terminal mutants in response to Copper and CDDP in Hek293T cell line. Fluorescence images showing localization of transfected Flag CTR1 (green) with plasma membrane marked with constitutive membrane marker Na^+^/K^+^ ATPase (Red) at Basal, copper(100µM) and CDDP (100µM). The Scale bar represents 10 microns. **(A)** Schematic diagram of membrane bound Ctr1 and targeted amino acids as mutants (Protter Software)**. (B)** Flag WT-CTR1 (*green*) localizes at plasma membrane at basal condition and endocytoses upon 100 μM Copper and CDDP treatment respectively. **(*C***) ΔH3-H6 CTR1 fraction localized at the plasma membrane under basal condition but exhibit distinct endocytosis in response to100 μM Copper but majority of the fraction stayed at the plasma membrane in response to CDDP. **(D)** Fraction of Δ30 Flag CTR1 shows significant colocalization with plasma membrane marker upon both Copper and CDDP treatment compared to basal. **(E)** The fraction of Flag-CTR1 Wt and respective N terminal mutants with membrane marker Na^+^/K^+^ ATPase demonstrated by a Box and Whisker plot with jitter points. (n>20) The box represents the 25 to 75th percentiles, and the median in the middle. The whiskers show the data points within the range of 1.5 × interquartile range (IQR) from the first and third quartile. ∗*p* < 0.05, ∗∗∗∗*p* < 0.0001; ns, not significant (nonparametric Mann–Whitney U test/Wilcoxon rank-sum test)

Histidine, owing to the presence of an imidazole group in its side chain, possesses a strong affinity for metal ions. To compare the potential role of the amino-terminal histidine residues of CTR1 in CDDP and copper uptake we utilized the Δ^3^HSHH^6^ deletion construct. Cells were transfected with ΔH-CTR1 and treated with 100µM CDDP or with 100µM copper. Under basal conditions, the mutant predominantly localized at the plasma membrane but underwent significant endocytosis upon copper exposure. In contrast, upon CDDP treatment the mutant Ctr1 displayed only mild endocytosis **(Fig.2C**). These observations led us to hypothesize that the ^3^H3SHH^6^ motif may coordinate with and sequester CDDP, though to a lesser extent than that of copper from the extracellular milieu, thereby facilitating its uptake through Ctr1.

Few previous studies have shown possible contribution of the distal amino-terminal harbouring the His motif and the M1 cluster in regulation of copper uptake and endocytosis of Ctr1 from the plasma membrane (5). We utilized the deletion construct, i.e., Δ30 CTR1 where first 30 amino acids of N-terminal harbouring the mentioned motifs of CTR1 is missing. Ctr1 endocytosis is not appreciable in either Copper and CDDP treated condition, though some endocytic vesicles were observed. Significant fraction of Δ30 CTR1 localized at plasma membrane marked with Na/K (**Fig.2D**). The comparative analysis of endocytosis based on colocalization between CTR1 and Na,K-ATPase is shown in **Fig.2E**. We further investigated individual roles of the amino-terminal Methionine clusters (M1: ^7^MGM^9)^ and (M2: ^40^MMMMPM^45^) in CDDP uptake and compared it with copper.

To determine role of Methionine clusters of CTR1 we utilized ΔM1 and ΔM2 constructs (illustrated in **Fig.3A**). In basal condition, both of these constructs exhibit localization on the plasma membrane. Interestingly ΔM1-CTR1 failed to endocytosed in response to copper; but in contrast upon CDDP exposure mutant protein significantly internalized from the plasma membrane. (**Fig.3B**). Similarly using ΔM2-CTR1 construct we observed minimal endocytosis of Ctr1 in response to CDDP as compared to basal condition. However, in copper treated condition, ΔM2-CTR1 exhibited moderate endocytosis (**Fig.3C**). These results indicate towards possible contribution of N-terminal Methionine residues, especially the second methionine cluster, in CDDP uptake while similar results were recorded for uptake of copper previously showed by Kar et al (5).

**Fig. 3.**
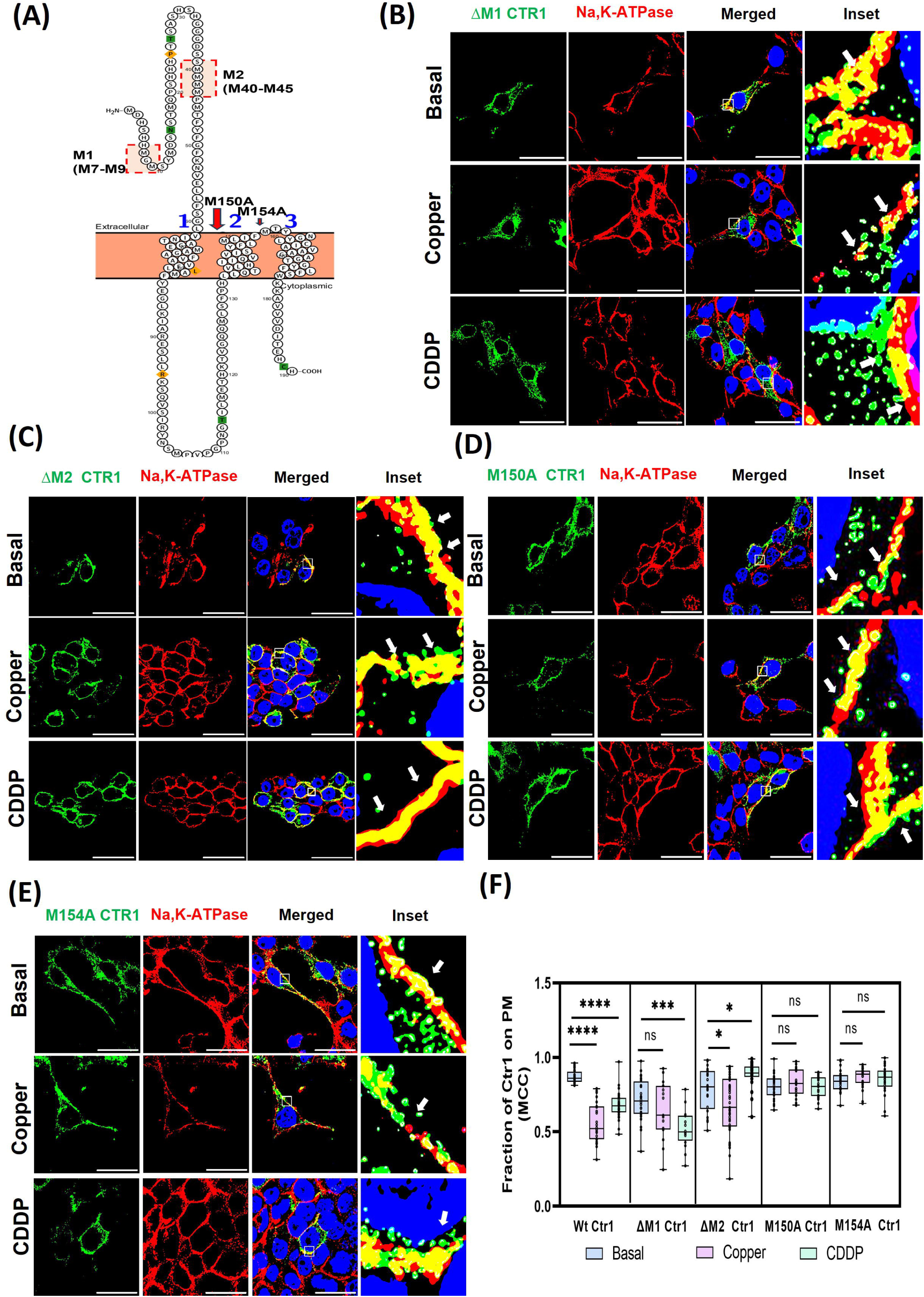
Methionine cluster on distal CTR1 N-terminal and in ion conduction pore on extracellular regulate CDDP uptake. Fluorescence images showing localization of transfected Flag CTR1 (green) mutants with plasma membrane marked with constitutive membrane marker Na^+^/K^+^ ATPase (Red) at Basal, copper(100µM) and CDDP (100µM). The Scale bar represents 10 microns. **(A)** Schematic diagram of membrane bound Ctr1 and targeted amino Met-mutants are drawn using Protter Software. **(B)** ΔM1 Ctr1 exhibits distinct membrane localization in both Basal and Copper treated condition and significantly endocytosed upon CDDP exposure. **(C)** Fraction of M2 Ctr1 (Green) showcase partial endocytosis in response to copper and CDDP and predominantly stayed at plasma membrane. **(D)** Copper and CDDP both failed to trigger endocytosis of M150A Ctr1 fraction from the plasma membrane. **(E)** M154A predominantly localize at the plasma membrane upon copper and CDDP exposure similar to basal condition. **(F)** The fraction of Flag-CTR1 Wt and respective Met-mutants with membrane marker Na^+^/K^+^ ATPase demonstrated using Box and Whisker plot with jitter points. (n>20) The box represents the 25 to 75th percentiles, and the median in the middle. The whiskers show the data points within the range of 1.5 × interquartile range (IQR) from the first and third quartile. ∗p < 0.05, ∗∗∗∗p < 0.0001; ns, not significant (nonparametric Mann–Whitney U test/Wilcoxon rank-sum test)

X-ray crystallography and mutagenesis studies have revealed that a central ion conduction pore formed between transmembrane helices 2 and 3 of Ctr1 is crucial for its channel-like function (32,37)The ^150^MXXM^154^ motif, located at the interface of these two transmembrane helices toward the extracellular surface, is evolutionarily conserved from lower eukaryotes to higher chordates and functions as a selectivity filter, facilitating the transfer of copper ions from the N-terminal domain through the channel to the C-terminal end of Ctr1 (12). To elucidate the role of the ^150^MXXM^154^ motif in CDDP uptake via Ctr1, we employed site-directed mutagenesis to substitute methionine residues at positions 150 and 154 with alanine. Under basal conditions, both mutants were localized at the plasma membrane and, interestingly, failed to undergo endocytosis upon exposure to either copper or CDDP (**Fig.3D & 3E).** Quantitative analysis of plasma membrane and endocytic fractions of Ctr1 under Basal, Copper, and CDDP-treated conditions for the N-terminal ΔM1 and ΔM2-CTR1, as well as the M150A and M154A mutants, is presented in **Fig.3F**.

To summarize, the extracellular amino-terminal His- and the second Met-rich motif (^40^MMMMPM^45^) of CTR1 are essential for CDDP recognition and uptake. Deletion or mutation of these residues, especially conserved ^150^MXXM^154^ motif, markedly reduces CDDP-induced CTR1 endocytosis thus highlighting CTR1 mediated CDDP import.

### CDDP and copper triggers distinct endocytic routes of CTR1

CTR1 exhibits endocytosis in response to both copper and CDDP, though in variable extents. We mapped the endocytic trafficking route of CTR1 in response to CDDP and compared to the pathway triggered by copper.

Unlike of copper induced CTR1 vesicles, the early endosomal marker, EEA1 do not show appreciable localization with CTR1 in response to 100µM and 50 µM CDDP but at low concentration, i.e., (25µM CDDP) a small fraction of CTR1 is observed on early endosomes (**Suppl fig S1A**).

Curnock et al showed the involvement of the VPS35, a subunit of the retromer complex in trafficking and recycling of Ctr1 between endosomal vesicles and plasma membrane, thereby influencing both copper homeostasis and cisplatin uptake (22). Contrastingly, we found that though copper induced significant localization of CTR1 in VPS35-positive compartments, CDDP did not induce similar localization of CTR1. (**Fig. 4B, S1B**).

**Fig. 4.**
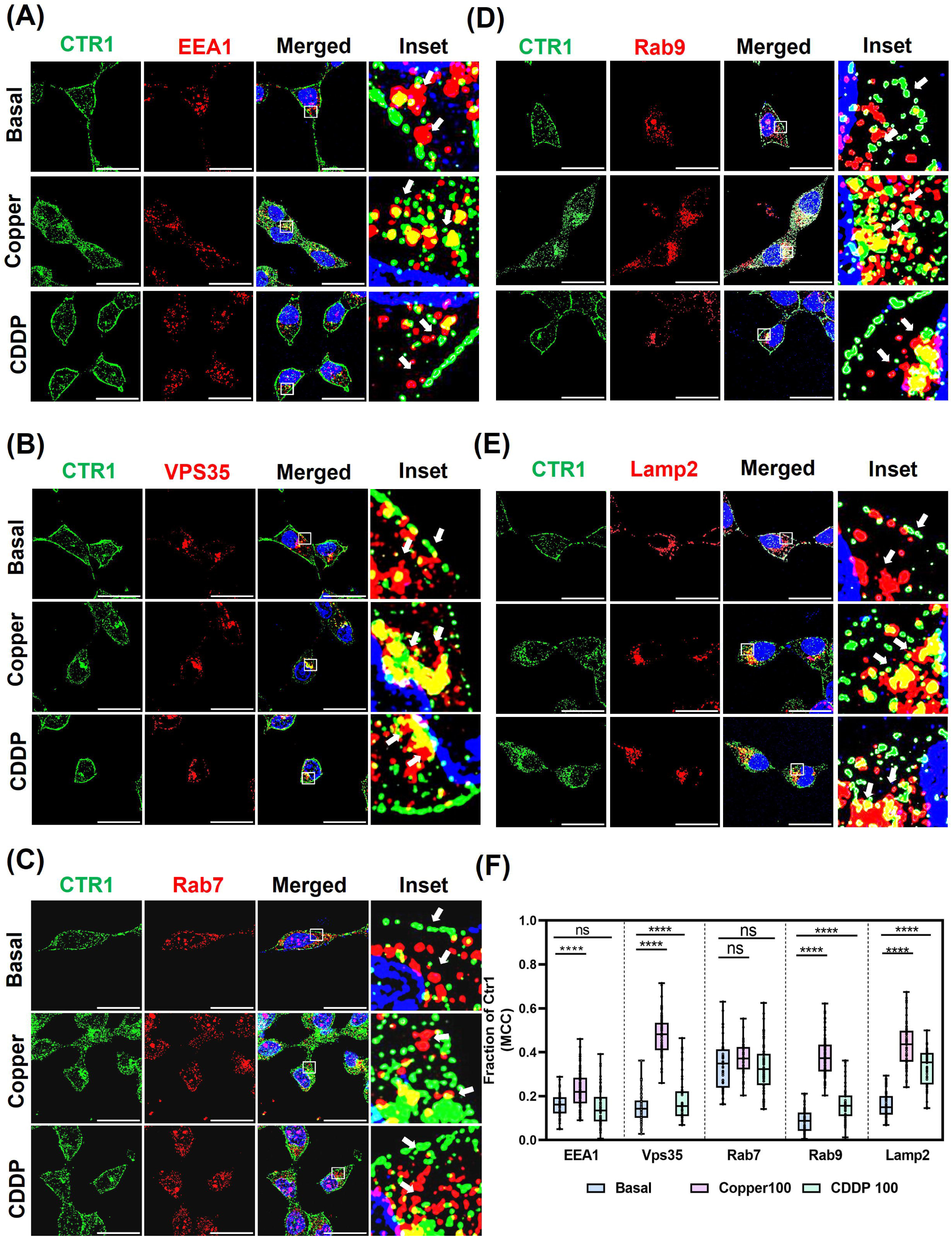
Copper and CDDP mediated endocytic sorting of Ctr1 in Hek293T cell line Fluorescence imaging showing fraction of CTR1 (green) localized with respective endocytic vesicles (red) while DAPI (blue) marks the nucleus at copper(100µM) and CDDP (100µM). The Scale bar represents 10 microns. **(A)** Immunofluorescence image showing fraction of Ctr1 (green) with early endosomal marker EEA1 (red) in Hek293T cell line. The marked area on the ‘merge’ panel is enlarged in ‘inset’. The arrowhead shows colocalization between Ctr1-EEA1. **(B)** Immunofluorescence image showing fraction of Ctr1 (green) with retromer controlled Recycling endosomal vesicle VPS35 (red) in HEK293T cell line. The marked area on the ‘merge’ panel is enlarged in ‘inset’. The arrowhead shows colocalization between Ctr1-VPS35. **(C)** Immunofluorescence image showing fraction of Ctr1 (green) with late endosomal marker Rab7 (red) in Hek293T cell line. The marked area on the ‘merge’ panel is enlarged in ‘inset’. The arrowhead shows colocalization between Ctr1-Rab7. **(D)** Immunofluorescence image showing fraction of Ctr1 (green) with retromer late endosomal marker Rab9 (red) in Hek293T cell line. The marked area on the ‘merge’ panel is enlarged in ‘inset’. The arrowhead shows colocalization between Ctr1-Rab9. **(E)** Immunofluorescence image showing fraction of Ctr1 (green) with lysosomal resident protein Lamp2 (Red) in Hek293T cell line. The marked area on the ‘merge’ panel is enlarged in ‘inset’. The arrowhead shows colocalization between Ctr1-Lamp2. **(F)** Quantitative analysis showing fraction of CTR1 residing in different endocytic vesicles upon 1 hr. treatment of copper (100 µM) and CDDP (100 µM) respectively graphically represented using a Box and Whisker plot with jitter points (n>50). The box represents the 25 to 75th percentiles, and the median in the middle. The whiskers show the data points within the range of 1.5 × interquartile range (IQR) from the first and third quartile. ∗*p* < 0.05, ∗∗∗∗*p* < 0.0001; ns, not significant.

Next, we tested localization of CTR1 on late endosomes upon CDDP treatment. Rab7 and Rab9 are both small GTPases in the Rab family that localizes at the late endosomes, but they have distinct roles in vesicular transport. Rab7 is a key regulator of late endosome and lysosome function, and transport to lysosomes, while Rab9 is primarily involved in retrograde transport from late endosomes back to the *trans*-Golgi network (TGN). Interestingly, in contrast to copper treatment, CTR1-Rab7 (late endosomal markers) exhibit very low colocalization at high CDDP treatments (**Fig.4C**). However, at lower concentration, i.e., 50µM CDDP, CTR1 shows slight increased colocalization on Rab7 vesicles, indication a possible pathway towards CTR1 degradation (**Fig.S1C**). Unlike with copper, CDDP did not induce localization of CTR1 in endosomes to Golgi recycling vesicles marked with Rab9 (**Fig.4D & S1D**). Consistent with CDDP triggering CTR towards late endosomal and further possibly towards lysosomal localization, at 100µM CDDP, lysosomal localization (marked by Lamp2) of CTR1 rises. We also recorded a gradual increase of CTR1 localization in lysosomes with increasing CDDP (25, 50 and 100µM) (**Fig.4E & S1E).** This indicates that CTR1 upon getting endocytosed owing to CDDP exposure, traverses through early-late endosomes towards the lysosomes for degradation. Alternately, the cell activates the lysosomal exocytosis pathway to export CDDP and to survive in high CDDP levels. Endosomal localization of CTR1 in copper and CDDP treatments have been quantitated in **Fig. 4F**.

### CDDP induces alteration in lysosome structure-function

CDDP induces lysosomal localization of CTR1. CDDP is released in the cell possibly from lysosome subsequent to which the drug is released in the intracellular milieu. Safaei and co-workers have shown that CDDP-resistant cells exhibit a significant reduction in lysosomal size, with only 40% of the lysosomal content compared to sensitive cells. Further, reduced expression of lysosome-associated proteins LAMP1 and LAMP2 is recorded in CDDP resistant cells (38).

Correlating with our observation of CDDP mediated late endosomal and lysosomal localization of Ctr1 and previous published studies, we tested whether CDDP induces alteration in lysosome physiology. We observed Cisplatin enhances Lysosomal activity by promoting lysosomal protease Cathepsins as analyzed by Magic Red assay; a Cathepsin B substrate which emits fluorescence depending upon Cathepsin B activity. (**Fig.5A & 5B**) We further observed from Particle analysis of Magic red puncta that moderate increase in Lysosome sizes occurs following 1hr treatment with CDDP (**Fig.5C**). Previous studies indicate that cathepsins exhibit maximal enzymatic activity in the acidic environment of lysosomes, generally maintained between pH 4.5 and 5.5.(39) Based on that we further tested the lysosomal acidity using Lysosensor Green, DND-189 and observed a significant increase in acidity of Lysosome using Flow cytometry which validate our observation with Magic Red following 1 hr. of Copper and CDDP treatment respectively (**Fig. 5D**). We did not record any appreciable change in lysosomal marker Lamp2 (**Fig.5E**, quantified in **Fig.5F**). Lysosomal biogenesis which signifies number of acidic compartments (Endo lysosome and Lysosome) remains unaltered when cells were probed with Lysotracker Red, DND-99ollowed by Flow cytometric analysis (**Fig. 5G**). This hints at a mechanism of CDDP mediated degradation of CTR1 and indicates an initial induction of Lysosomal activity following 1hr exposure to CDDP.

**Fig. 5.**
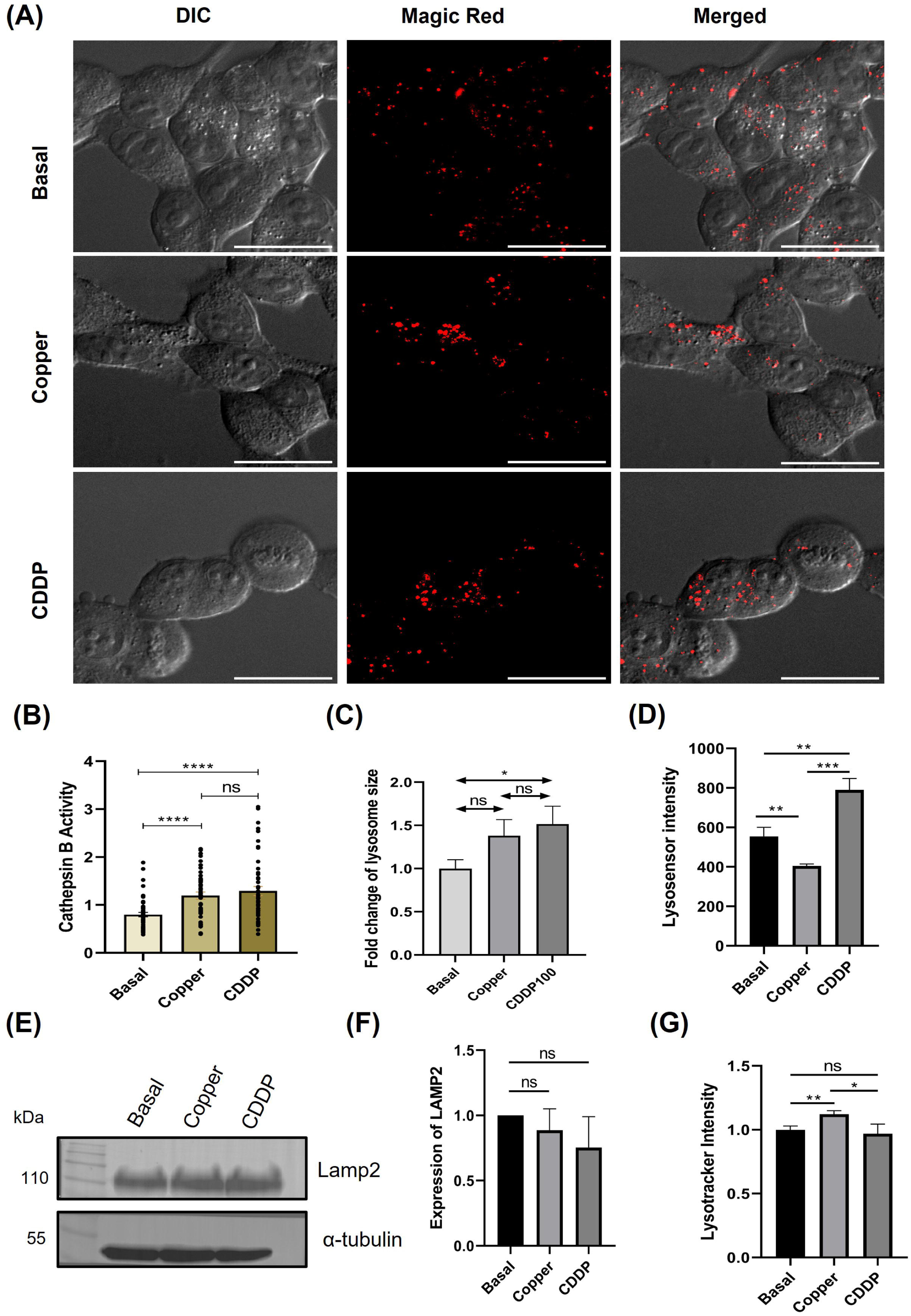
Copper and CDDP regulate lysosomes distinctly. **(A)** Magic red, a substrate of Cathepsin B which signifies lysosomal activity showed via immunofluorescence images in Hek293T cell line. Red dots indicate cathepsin positive lysosomes in Copper and CDDP treated condition. **(B)** Cathepsin B activity measured using Corrected total cell fluorescence (CTCF) analysis and plotted **(C)** Size of cathepsin positive lysosomes analyzed and plotted. **(D)** Lysosensor Green, dye measuring change in lysosomal P.H used to measure acidic characteristics of Lysosome via Flow cytometry(n=3). **(E)** Expression of Lamp2, a lysosomal membrane resident glycoprotein associate with lysosomal number and function showed by Western blot. **(F)** Abundance of Lamp2 in Hek293T cells analyzed using Non-Parametric T test and plotted. (n=3) **(G)** Number of acidic lysosomal compartments measured using Lysotracker via Flow cytometry analyzed and plotted**. (n=3)**

### CDDP selectively alters some cuproproteins significantly

CTR1 is the sole high affinity copper importer in mammalian cells (11). Once uptaken, free copper is quickly sequestered by chaperones, ATOX1, CCS and COX17 being the major cupro-chaperones. Excess copper is exported by the copper-ATPases, ATP7A and ATP7B. Together, the synchronized functions of copper chaperones, importers and copper exporting ATPases keeps the intracellular copper levels in tight regulation. Though it has been established that CDDP employs CTR1 to gain entry into cells, it is still unclear whether CDDP also uses the copper transport pathway that involves proteins like the copper chaperone for SOD1, CCS. Safaei and colleagues showed that the metallochaperone, ATOX1 helps cisplatin enter cells by promoting Ctr1 internalization and ubiquitination, which increases drug sensitivity, while the loss of ATOX1 lowers cisplatin uptake (40).

To examine the impact of CDDP on copper homeostasis pathways, we analyzed the expression of various cuproproteins using immunoblotting. Here, we extended our analysis to evaluate the effect of a range of CDDP concentrations, selected based on MTT survival assays in HeLa and HEK293 cells (**Fig.1A & 1B**), on the expression of multiple cuproproteins in HEK293 cells.

The immunoblot analysis in **Fig. 6A**, along with its corresponding quantification in **Fig. 6B**, revealed that CTR1 levels increased slightly though significantly with rising copper concentrations (1-hr treatment), whereas they gradually decreased with increasing concentrations of CDDP. We specifically quantified the glycosylated fraction of CTR1 (35 kDa), given its localization at the plasma membrane.

**Fig. 6.**
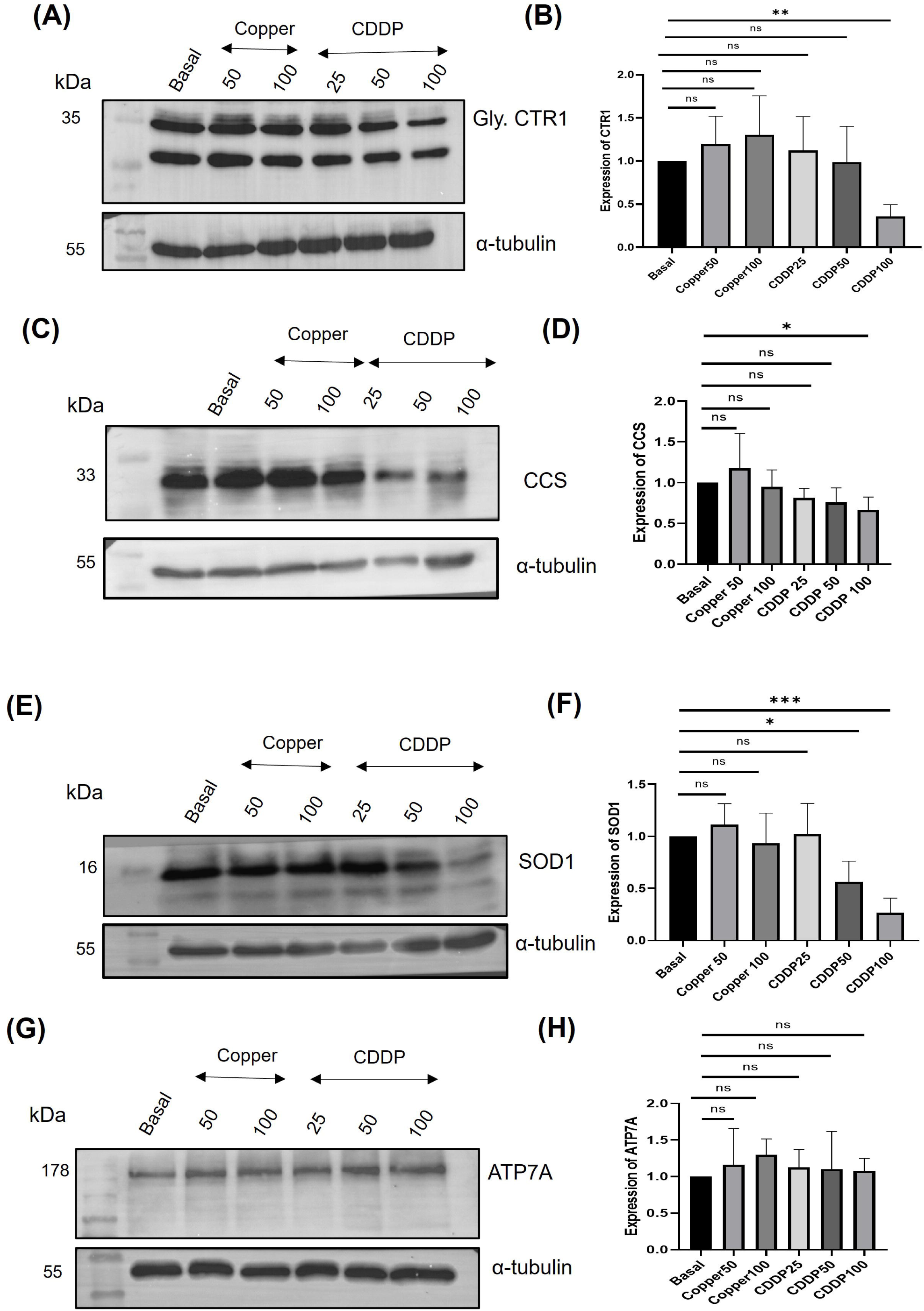
Copper and CDDP regulates levels of cuproproteins differently. **(A)** Immunoblot of CTR1 following 1 hr exposure to indicated conc. of copper and CDDP. Fold change of Glycosylated fraction of CTR1(Arrowed) abundance normalized against housekeeping control α-tubulin has been mentioned and plotted in **6B**.(n=3) **(C)** Immunoblot of CCS following 1 hr exposure to indicated conc. of copper and CDDP. Fold change of CCS abundance normalized against housekeeping control α-tubulin has been mentioned and plotted in **6D.** (n=3) **(E)** Immunoblot of SOD1 following 1 hr exposure to indicated conc. of copper and CDDP. Fold change of SOD1 abundance normalized against housekeeping control α-tubulin has been mentioned and plotted in **6F**.(n=3) **(G)** Immunoblot of ATP7A following 1 hr exposure to indicated conc. of copper and CDDP. Fold change of ATP7A abundance normalized against housekeeping control α-tubulin has been and plotted in **6H**.(n=3). Error bars represent mean□±□SD of values calculated from three independent experiments. Asterisks indicate values that are significantly different from samples. ^∗^*p*□≤□0.05, ^∗∗^*p*□≤□0.01, ^∗∗∗^*p*□≤□0.001, ns; (Student’s *t* test).

CCS and SOD1, the cytosolic copper chaperones, showed a decline in expression with increasing concentrations of CDDP, whereas their levels are unchanged in response to 50 µM and 100 µM copper treatment (**Fig 6C, 6D & 6E, 6F).**

In contrast, the expression levels of other copper transport pathway proteins remained largely unchanged. Notably, ATP7A, the copper exporting-ATPase, also implicated in CDDP efflux and drug resistance (41) did not show a significant increase in expression following CDDP exposure (**Fig. 6G, 6H).**

These findings suggest that cisplatin and copper interact differently with downstream cuproproteins, indicating distinct modes of action after entering the cell through CTR1, which likely serves as the primary entry route.

## Discussion

Copper is an essential trace element that functions as a critical cofactor for a wide range of metalloenzymes within cells. Copper exists primarily in two oxidation states, Cu(I) and Cu (II). Of these, Cu(I) is generally considered the biologically active form. CTR1 is a sole protein widely distributed through the plasma membrane of eukaryotic life forms; responsible for importing copper into cells. The pharmacological significance of CTR1 was first reported by Ishida et al. demonstrating that CTR1 mediates the cellular accumulation of the anticancer drug CDDP (23).

Previous studies have elucidated the structural basis of copper import, demonstrating critical roles for the N-terminal domain, transmembrane segments, and C-terminal region of CTR1, as well as its endocytosis into subcellular compartments to prevent copper overload. Although the role of CTR1 in mediating the cellular uptake of the anticancer drug CDDP is well documented, the precise mechanism of CDDP transport through CTR1 and the fate of CTR1 following drug exposure remain unresolved.

In this study, utilizing mutational approach and immunofluorescence microscopy, we demonstrated the involvement of the amino-terminal domain of CTR1 in CDDP uptake. Furthermore, we show the detailed sorting map of CTR1’s intracellular trafficking in response to varying concentrations of CDDP and compare it with the sorting pathway utilized by CTR1 in response to copper. By tracking CTR1 trafficking, we observed that a substantial fraction of the protein localized to endolysosomal and lysosomal compartments after 1 h of CDDP exposure. Interestingly, in response to CDDP, though CTR1 does not traverse the early endosomes marked by EEA1, low doses of CDDP (25µM) makes it retained in the VPS35 positive compartments. Higher levels of CDDP (50µM and 100µM), reduces this colocalization between VPS35 and CTR1. We can extrapolate that at low levels of CDDP, the retromer complex comprising of VPS26, VPS29, VPS35 and the SNX proteins can steer the trafficking direction of CTR1 back towards the plasma membrane. However, it is also possible that the retromer complex regulates sorting of CTR1 towards the endolysosomal or lysosomal compartments. We have previously demonstrated that VPS35 sorts the copper ATPase, ATP7B at the endolysosomal compartments and directs it toward active lysosomes in response to high copper (42). It is evident that with increasing levels of CDDP (25µM, 50µM and 100µM), CTR1 localizes at the endolysosomal and the lysosomal compartments marked by Rab7 and LAMP2 respectively. This points towards the fact that CDDP release in the cell possibly if facilitated by lysosomal functioning. To further investigate to this response, we examined the effects of CDDP on lysosomal physiology.

Initial examination of cell survivability using cytotoxicity and yeast growth assays we highlighted the competition between copper metabolism and CDDP uptake. While copper supplementation increased their survival, copper-chelator treatment further enhanced CDDP sensitivity and reduced viability. These findings indicate that CDDP’s effect is closely tied to cellular copper levels, with CTR1 and copper-regulating pathways influencing drug response. This conclusion is further supported by experiments using CTR1 wild-type and knockout yeast strains, where the knockout cells displayed a higher survival rate compared to wild-type cells when exposed to CDDP.

We further asked fate of plasma membrane localized CTR1 in response to CDDP? Clifford et al reported dynamic internalization of CTR1 in response to copper and further demonstrated the fate of endocytosed CTR1 at different endosomes in response to copper (43). Similarly, upon increasing CDDP concentrations up to 100 µM, we observed endocytosis of CTR1 from the plasma membrane, resembling its behavior in the presence of copper. This internalization may represent a prominent mechanism of resistance to CDDP.

Studies from our group previously demonstrated role of -Met, -Asp and -His residues of N-terminal domain of CTR1 to reduce Cu (II) to Cu(I) and subsequent import of copper to cytosol (5). Failure of the Δ30 mutant of CTR1 upon copper or CDDP treatment highlights the importance of the first Met motif and the His motif. Interestingly, a different pattern was observed with first methionine cluster mutants of CTR1, where the protein failed to undergo endocytosis in response to copper but displayed partial endocytosis at low to moderate levels upon CDDP exposure. We establish the extracellular N-terminal domain as a key CTR1 domain playing critical role in CDDP uptake.

Our findings indicate that cisplatin and copper exert distinct effects on downstream cuproproteins despite sharing CTR1 as a common entry route. While copper upregulated the expression of CCS and SOD1, cisplatin reduced their levels, and ATP7B remained largely unaffected under both conditions. It is possible that once uptaken, CDDP is sequestered by CCS-SOD pathway in addition to being bound to ATOX1 and subsequently by ATP7A (41,44). This differential regulation highlights that cisplatin does not simply mimic copper but instead hijacks the copper transport pathway in a unique manner, potentially contributing to its cytotoxic activity and resistance mechanisms.

To summarize we propose an involvement of N-terminal of CTR1 in CDDP import and eventual endocytosis of CTR1. However, we observe that the mechanism of uptake and sorting pathways of CTR1 in response to copper and CDDP are not exactly the same. This differential regulation highlights that cisplatin does not simply mimic copper but instead hijacks the copper transport pathway in a unique manner, potentially contributing to its cytotoxic activity and resistance mechanisms. The proposed model is illustrated in **Fig. 7**. It will be important to investigate the role of other amino acid residues of CTR1 in cisplatin uptake and to study in detail the cellular effects of chronic cisplatin exposure on organelles such as lysosomes, mitochondria, and endosome. As direct interplay between CDDP and Copper for its import, these studies will provide deeper insights into how a drug can differentially affect pathways involving an essential micronutrient like copper. The findings will provide a better mechanistic view of CDDP drug-resistance during prolonged chemotherapeutic treatment.

**Fig. 7.**
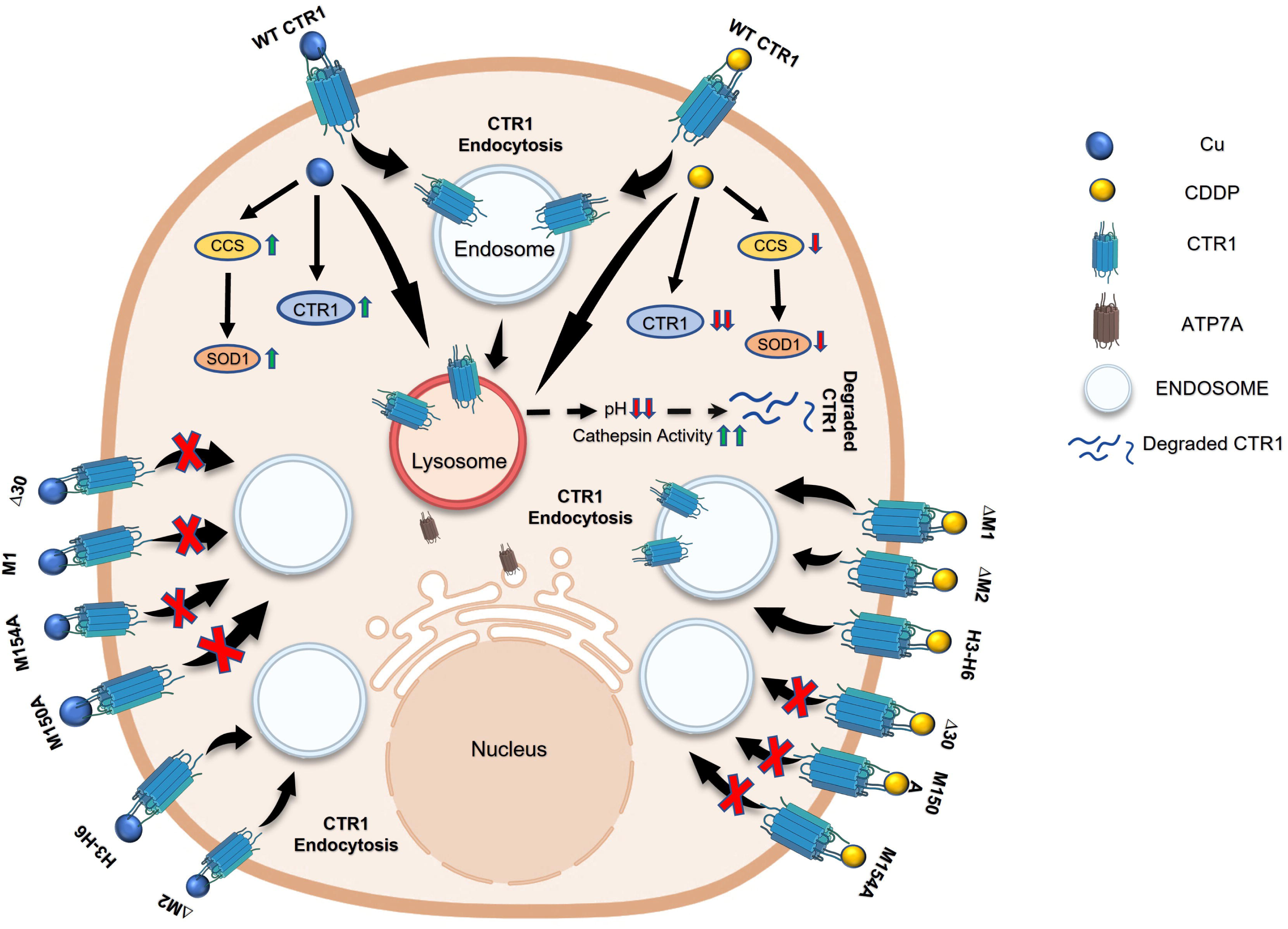
Schematic illustration showing how both Copper (blue dot) and Cisplatin (CDDP; yellow dot) trigger CTR1 endocytosis from the plasma membrane of unpolarized embryonic kidney HEK293T cells. Following internalization, CTR1 traffics through the endosomal pathway to the lysosome upon both Copper and CDDP treatment. Mutant CTR1, ΔH3–H6, ΔM1, Δ30, ΔM2, M150A, and M154A exhibit differential internalization responses to copper or CDDP. Arrows indicate endocytic trafficking of CTR1 to intracellular vesicles, while red crosses denote blocking of endocytosis. Upon uptake through CTR1, Copper and CDDP differentially modulate cellular cuproprotein responses—Copper increases CCS and SOD1 levels, whereas CDDP reduces their expression. Unlike Copper, CDDP primarily affects lysosomal function by decreasing lysosomal pH and enhancing cathepsin activity.

## Materials and Methods

### Cell lines, cell culture and transfection

Non-cancerous epithelial HEK293 cells and cancerous cervical HeLa cells were maintained in Dulbecco’s modified Eagle’s medium (#D6429; Sigma) supplemented with 10% fetal bovine serum (FBS, Gibco #26140079) and 1% penicillin–streptomycin (Gibco #15140122) in 5% CO_2_ incubator at 37°C. FlagCtr1-Wt and N-terminal mutations plasmids were used and transfection were performed using the Jet PRIME transfection reagent (Polyplus, #114-07) as per the manufacturer’s instructions and protocols were followed from previous studies using same plasmids of our Lab.(**Kar et al.**)

### Copper, CDDP and other treatments

A 10 mM stock solution of copper chloride (SRL #92315), prepared by dissolving it in water, was used as the source for a 50 µM and 100 µM copper treatment to study trafficking and immunoblot assays. Similarly, a 10 mM stock solution of Cisplatin (CDDP, TOCRIS #2251), prepared by dissolving it in DMF (N,N-Dimethylformamide, Sigma #227056), was used for 25 µM,50 µM and100 µM cisplatin treatments, to investigate the same experiments as those conducted with copper. Further, 25 µM TTM (Tetra thiomolybdate, Sigma #323446) dissolved in DMSO (dimethyl sulfoxide, Sigma #276855) was used as copper chelator prior to CDDP treatment. Copper, CDDP and TTM treatments were applied for 1 H. Small molecule like Bafilomycin A1 (lysosomal inhibitor, Sigma;#SML1661) was used for 1hr before 1hr of Copper and CDDP treatment respectively.

### Cell viability assay

The colorimetric MTT reduction assay was conducted to assess cell viability post Cu/TTM treatment along with CDDP. HeLa or HEK293 cells cultured on 96-well plates were pre-treated with Cu or TTM for 1 H followed by increasing concentrations of CDDP for further 1 H. The cells were then incubated with 1 mg/ml MTT solution (Sigma #M2128) at 37 °C for 3 h and formazan grains formed were dissolved in DMSO. The optical density (OD) of dissolved formazan grain was measured at 595 nm by a Multilabel Plate Reader (GeneX). Cell viability was expressed as a percentage relative to the value in untreated controls (100%).

### Immunofluorescence and microscopy

Cells were cultured in round coverslips placed in 24 well plates and all the solutions mentioned above were treated once the cells reached 70-80% confluency. Cells were washed with ice cold PBS after 1 H treatment. Flag Ctr1 transfected cells were fixed using 2% Paraformaldehyde for 20 mins followed by PFA quenching using 50 Mm NH_4_cl for 20mins where fixation with aceto-methanol (acetone-methanol 1:1) for 10 min at room temperature performed to the cells probe with endogenous Rabbit Ctr1 and other trafficking Antibodies. Next blocking was performed in 3% bovine serum albumin (BSA, SRL #85171) in PBSS (0.075% saponin in PBS) for 2 H. Primary antibody (in 1% BSA in PBSS) incubation was conducted for 2 H at room temperature followed by respective secondary antibody incubation for 1 H. Finally, coverslips were mounted on glass slides using SIGMA Vectashield^TM^ with DAPI mounting media (Sigma #F6057). All images were captured using a Leica SP8 confocal system with a 63x oil immersion objective (NA 1.4) and processed through deconvolution with Leica Lighting software.

The primary antibodies utilized in immunofluorescence studies are as follows: Rabbit anti-FLAG M2 (1:600,CST #14793), Rabbit Ctr1(1:800,# ab129067 Abcam), Mouse Na/K ATPase (1:600,#MA3-929 Invitrogen), Mouse VPS35 (1:500,#sc-374372, Santa Cruz Biotechnology), Mouse Rab7 (1:50,#sc-376362, Santa Cruz Biotechnology), Mouse Lamp2 (1:150, #H4B4 DSHB), Mouse Rab9 (1:100, #MA3-06 Invitrogen), Mouse EEA1(1:100 #610457, 1:400; BD Biosciences).The Secondary antibodies utilized in immunofluorescence studies are as follows: Donkey anti-Rabbit IgG (H + L) Alexa Fluor 488 (1:2000, #A-21206 Invitrogen), Donkey anti-Mouse Alexa 555 (#A-32773, 1:2000; Invitrogen).

### Incubation with Cathepsin B Substrate (Magic Red assay)

To label endocytic organelles in which cathepsin B was catalytically active, HepG2 cells were incubated with the Magic Red substrate from ImmunoChemistry Technologies, as per manufacturer’s instruction.

### Yeast growth rate assay

*Saccharomyeces cerevisiae* BY4742 (WT) strain (MATα his3Δ1 leu2Δ0 lys2Δ0 ura3Δ0) and *S. cerevisiae* strain carrying any CTR1 deletion (*yCTR1*) in the BY4742 background purchased from Euroscarf (Oberursel) were used. YPD (yeast extract, peptone, and dextrose) medium was used to grow both WT and deletion strains, and growth were monitored every hour in presence and absence of CDDP. In order to measure the growth rates of the yeast strain, the yeast cultures grown up to saturation at a density of OD_600_ 0.2 were further diluted and grown at 30 °C incubator for around 24 h, and growth density was measured spectroscopically after 0, 1, 3, 6, 9, 10,12, 18, and 24 h. Growth curves of yeasts expressing WT and mutant hCTR1 in YPEG culture media were generated over the 24-h period. Yeast growth rates (ΔOD_600_/hour) were calculated from linear exponential growth phase between 3 and 12 H time points.

### Immunoblot assay

Cells were grown on 60 mm dishes and treated with copper and CDDP treatments followed by protein isolation. Cell pellet thus isolated was dissolved in 100 μl of RIPA lysis buffer (10=mM Tris-Cl pH 8.0, 1=mM EDTA, 0.5=mM EGTA, 1.0% Triton X-100, 0.1% sodium deoxycholate, 0.1% SDS, 140=mM NaCl, 1X protease inhibitor cocktail, 1=mM phenylmethylsulphonyl fluoride (PMSF) and incubated on ice for 1 H. The lysates were then sonicated (four pulses; 5 s on phase and 30 s off phase; amplitude 100 mA) with a probe sonicator. Bradford was used for protein quantitation and 25 μg protein was loaded in each well. The Protein samples were prepared by adding 4X NuPAGE loading buffer (#NP0007; Invitrogen) to a final concentration of 1X and run on SDS-PAGE (10-12%) gels to separate proteins according to molecular mass. This was followed by semi-dry transfer (Bio-Rad Trans-Blot SD Cell) of proteins onto nitrocellulose membrane (#IPVH00010; Millipore). Following transfer, the membrane was blocked with 3% skimmed milk/BSA in 1X Tris-buffered saline (TBS) buffer pH 7.5 for 2 H at RT with mild shaking. Membrane containing proteins were incubated with Primary antibody was done overnight at 4 °C then washed with 1X TBST.HRP-conjugated respective secondary antibody incubation for 2 H at RT was performed followed by 3 TBST and 2 TBS washes. The chemiluminescent signal was developed by Clarity Max Western ECL substrate (BioRad #1705062) through chemiluminescence by Chemi Doc (BioRad).

### Flow cytometry assay

HEK293T cells seeded on 12-well plates were treated with 100 µM of Copper and CDDP for 1 hr at 70-80% confluency. After treatment cells were washed with 1xdPBS.Following the washes cells are incubated with 50nM Lysotracker Red DND99 (#L7528; Thermo) and 1µM Lysosensor Green DND-189 (#L7535; Thermo) for 30 mins.Cells were washed with 1XDPBS and kept in 50 µL trypsin for 2 min.After that cells were suspended in 350 µL of FACS buffer (1X DPBS, 2% FBS, 25=mM HEPES, 2=mM EDTA) and analysed using Flow cytometry BD LSRFortessa (BD Biosciences).Signals from Ten thousand cells were quantitate for each condition. FCS Express software (version 7.14.0020) was used for data analysis. Mean intensity values were used for quantification.

### Image analysis and statistics

Images were analysed in batches using ImageJ. For the colocalization study, the Colocalization Finder plugin was utilized. Regions of interest (ROIs) were manually drawn on the optimal z-stack for each cell. Manders’ colocalization coefficient (MCC)(45) and Pearson correlation coefficient was employed to quantify colocalization(46). ImageJ macro codes for image analysis was used in R v-42.1 (47) In the box plots, the boxes indicate the interquartile range (25th–75th percentiles), with the horizontal line representing the median. Whiskers extend to data points falling within 1.5 times the interquartile range from the first and third quartiles. Statistical comparisons between unpaired datasets were performed using non-parametric tests (Mann–Whitney U test/Wilcoxon rank-sum test). Significance levels were denoted as follows: *p=<=0.05; **p=<=0.01; ****p=<=0.0001; ns, not significant. All experiments performed independently for at least three times. Graphs were generated using GraphPad Prism software (version 9.4).

## Acknowledgement

This work was supported by DBT-Wellcome Trust India Alliance Fellowship (IA/I/16/1/502369) and Core Research Grant (CRG/2021/002150) from SERB, Department of Science and Technology (DST), Government of India STARS-2 Grant (2023-0210) from Ministry of Education, Govt. of India and IISER K intramural funding to A.G. M.C. was supported by pre-doctoral fellowship from University Grants Commision, Govt. of India. S.K was supported by pre-doctoral fellowship from Council of Scientific and Industrial Research, Govt. of India. A.P. was supported by National Postdoctoral Fellowship from SERB, Govt. of India.

## Author contributions

A.G conceptualized and designed the study, drafted paper and acquired funding for the study, M.C conducted experiments and analyzed data, A.P conducted experiments and analyzed data, S.K conducted experiments

## Supplementary figures

**Figure S1.**
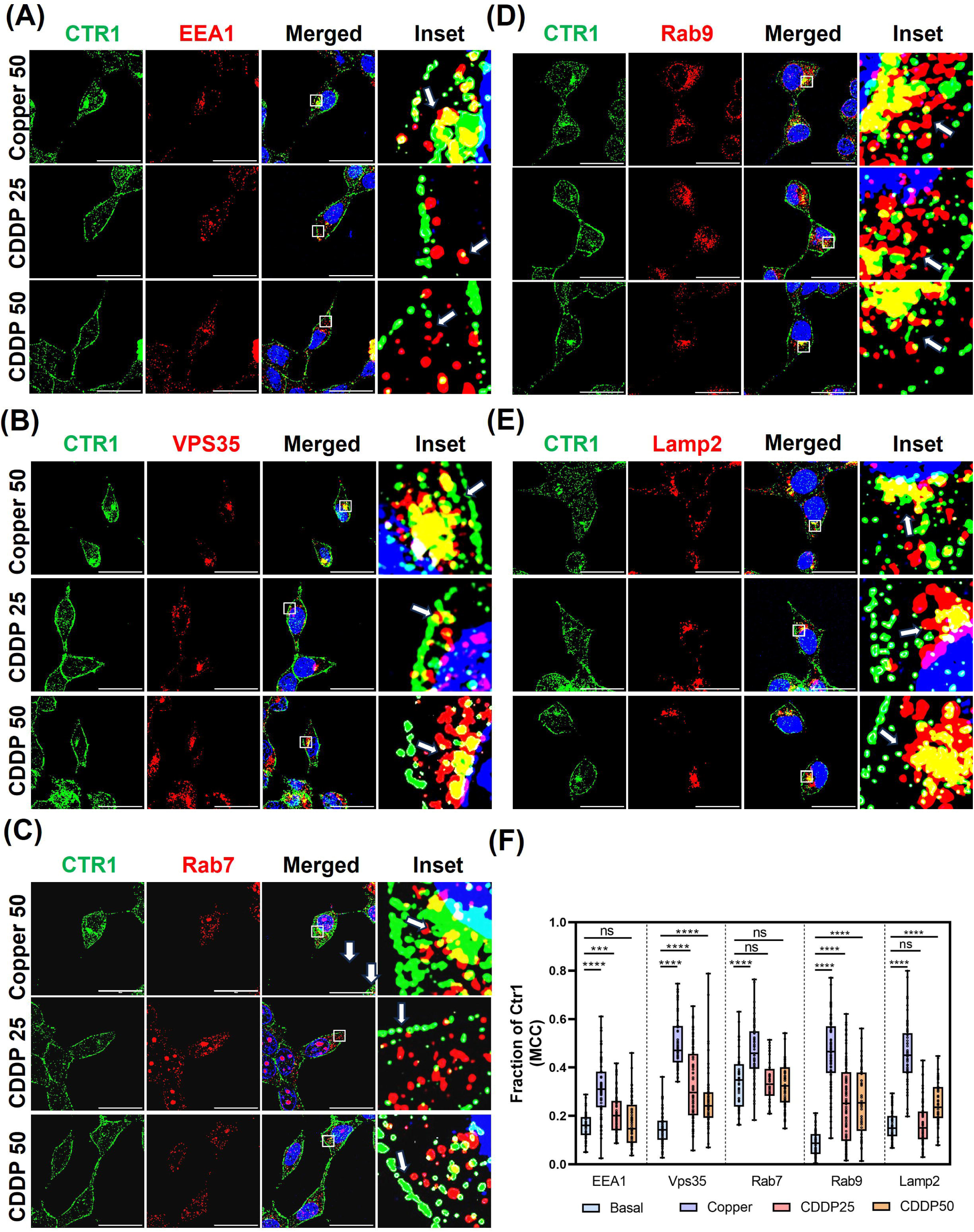
Fluorescence imaging showing fraction of CTR1 (green) localized with respective endocytic vesicles (red) while DAPI (blue) marks the nucleus at copper (50 µM) and CDDP (25µM & 50µM). The Scale bar represents 10 microns. **(A)** Immunofluorescence image showing fraction of Ctr1 (green) with early endosomal marker EEA1 (red) in Hek293T cell line. The marked area on the ‘merge’ panel is enlarged in ‘inset’. The arrowhead shows colocalization between Ctr1-EEA1. **(B)** Immunofluorescence image showing fraction of Ctr1 (green) with retromer controlled Recycling endosomal vesicle VPS35 (red) in Hek293T cell line. The marked area on the ‘merge’ panel is enlarged in ‘inset’. The arrowhead shows colocalization between Ctr1-VPS35. **(C)** Immunofluorescence image showing fraction of Ctr1 (green) with late endosomal marker Rab7 (red) in Hek293T cell line. The marked area on the ‘merge’ panel is enlarged in ‘inset’. The arrowhead shows colocalization between Ctr1-Rab7. **(D)** Immunofluorescence image showing fraction of Ctr1 (green) with retromer late endosomal marker Rab9 (red) in Hek293T cell line. The marked area on the ‘merge’ panel is enlarged in ‘inset’. The arrowhead shows colocalization between Ctr1-Rab9. **(E)** Immunofluorescence image showing fraction of Ctr1 (green) with lysosomal resident protein Lamp2 (Red) in Hek293T cell line. The marked area on the ‘merge’ panel is enlarged in ‘inset’. The arrowhead shows colocalization between Ctr1-Lamp2. **(F)** Quantitative analysis showing fraction of CTR1 residing in different endocytic vesicles upon 1 hr. treatment of copper (100 µM) and CDDP (100 µM) respectively graphically represented using a Box and Whisker plot with jitter points (n>50). The box represents the 25 to 75th percentiles, and the median in the middle. The whiskers show the data points within the range of 1.5 × interquartile range (IQR) from the first and third quartile. ∗*p* < 0.05, ∗∗∗∗*p* < 0.0001; ns, not significant.

